# Improved Degradome Sequencing Protocol via Reagent Recycling from sRNAseq Library Preparations

**DOI:** 10.1101/2024.08.04.606535

**Authors:** Marta Puchta-Jasińska, Jolanta Groszyk, Maja Boczkowska

**Affiliations:** Plant Breeding and Acclimatization Institute, Radzików, 05-870 Błonie, Poland

**Author notes:** Correspondence: Marta Puchta-Jasińska.

**Keywords:** degradome, cost effective, protocol, miRNA, target genes

## Abstract

**Background:** One of the key elements in the analysis of gene expression and its post-translational regulation is miRNAs. Degradome-seq analyses are performed to analyze the cleavage of target RNAs in the transcriptome. In this work, an improved library preparation protocol for degradome sequencing is presented. The developed protocol improves the efficiency of library preparation in degradome-seq analysis used to identify microRNA targets, reduces the time of library preparation and lowers the cost of purchasing reagents..

**Results:** The aim of this study was the development of an efficient protocol for the construction of degradome sequencing libraries using residual reagents from the sRNA-seq library preparation kit. To this end, modified primers and adaptors were designed. The library purification step based on automated electrophoresis and high-resolution agarose was modified and optimized in the presented protocol. Size standards of 60 and 65 bp were developed. They were prepared for precise band excision from the gel. Cloning to plasmid and sequencing of the inserted fragment, i.e., a fragment from the degradome library, verified the correctness of the library preparation using the developed protocol.

**Conclusion:** The developed protocol allowed the construction and sequencing of degradome libraries even from RNA samples with low RIN. It significantly reduces the cost of library construction. This is due to the use of residues from the sRNA-seq library kit. The precision of the excised fragment after electrophoresis performed during the procedure to isolate fragments of the correct length is significantly improved by the use of additional size markers. Compared to previously used methods, optimizing the purification method of degradom-seq libraries allowed to increase the yield of fragments obtained. Notably, the time required for the entire library preparation protocol does not exceed three days, also a significant time savings.

## Background

In plants, microRNAs (miRNAs) primarily function by cleaving of target messenger RNAs (mRNAs) molecules, which have near-perfect complementarity at the matching sites [1]. Computational methods based on genome-wide searches for targets for miRNAs, which rely on base pairing and assessment of miRNA: target interactions have been shown to yield a high percentage of false-positive predictions [2]. The pairing of miRNAs with target genes is a complex process influenced by numerous factors, including expression patterns over time. Therefore, experimental verification of targets is crucial step in the research process [3]. In 2008, three methods for the verification of miRNA targets based on next-generation sequencing were published i.e. parallel analysis of RNA ends (PARE) [4], degradome sequencing (degradome-seq) [5] and genome-wide mapping of uncapped and cleaved transcripts (GMUCT)[6]. The aforementioned methods are founded upon the capture of a sequence devoid of a cap at the 5’ side of a cut 3’ mRNA that has been ligated with an adapter [5]. Degradome-seq represents a high-throughput sequencing method that draws inspiration from a modified 5’-rapid amplification of cDNA ends (5’-RACE) approach. Degradome sequencing enables the identification of target genes that are subjected to cleavage by miRNAs. The methodology presented here constitutes a modification of the protocol originally described by Lin at el. (2019)[7]. method facilitates the preparation of libraries in a more streamlined and efficient manner, while also enabling the utilization of residual components from the NebNext small RNA Library Prep Set for Illumina (New England Biolabs) following the preparation of miRNA sequencing libraries. Degradome profiles can provide substantial insight into the intricacies of RNA processing [7]. the method generates a fixed fragment length library through enzymatic restriction using Mme I, which cleaves 20 bp 3′ of the recognition site included in the 5′ cDNA adaptor. Subsequently, 3’ dsDNA adapters are ligated to the digestion products, and the fragments are then subjected to PCR amplification, purification, and appropriate sequencing[1]. The analysis of networks regulating miRNA-mediated gene expression based on degradome sequencing data presents a superior approach to the computational (*in silico*) prediction of miRNA targets. Additionally, interactions between siRNAs and alterations in gene expression can also be monitored using degradome sequencing data [8]. Recently, studies have also demonstrated the potential of such data in the field of RNA research. For example, it can be used to map endoribonucleolytic cleavage sites in vivo, identify conserved motifs at the ends of 5’ naked RNA fragments and the search for regions associated with stacked ribosomes (or other RNA-binding proteins) on transcripts[8]. The rate of false-positive predictions of miRNA binding sites and the size of the search space for miRNA target sites were significantly reduced by the application of CLIP-Seq and Degradome-seq methods. A crucial aspect of elucidating the biological function of small regulatory RNAs (sRNAs) is the identification and validation of their targets. The majority of computational tools utilized for the prediction of sRNA targets in plants (and animals) employ techniques that seek to identify complementarity between an sRNA sequence and a potential target sequence[9].

## Materials

### Reagents

- TRI Reagent (Sigma-Aldrich, Saint Louis, MO, USA; T9424)
- 1-bromo-3-chloropropane (Sigma-Aldrich, Saint Louis, MO, USA; B9673)
- 2-propanol (POCH, Gliwice, Poland; 751503420)
- 99,8% ethanol A.C.S. (Chempur, Piekary Śląskie, Poland; 113964800)
- Molecular biology water (A&A Biotechnology, Gdynia, Poland; 003-075) water non-treated withDEPC
- KoAc (Sigma-Aldrich, Saint Louis, MO, USA; P1190)
- Dynabeads Purification Kit for mRNA (Invitrogen, Carlsbad, CA, USA; 61006)
- NebNext small RNA Library Prep Set for Illumina (New England Biolabs, Ipswich, MA, USA; E7300S)
- Ribolock (40 U/µL) (ThermoFisher Scientific, Waltham, MA, USA; E00381)
- Maxima H Minus First Strand cDNA Synthesis Kit (200 U) (ThermoFisher Scientific, Waltham, MA, USA; K1652)
- AmPure XP for PCR Purification (Beckman Coulter Life Sciences; Indianapolis, United States; A63880)
- SeqAmp DNA Polymerase (Takara Bio, Otsu, Japan; 638504)
- Customer primers (10 µM)
- Phix Control Sequencing v3 (Illumina, San Diego, CA, USA; FC-110-3001)
- Illumina MiSeq, Reagent Kit v3 (50-cycles) cartridge (Illumina, San Diego,CA, USA; MS-102-2001)

NaOH

- Sodium acetate (Sigma-Aldrich, Saint Louis, MO, USA; P1190)
- Glycogen (ThermoFisher Scientific, Waltham, MA, USA; R0561)
- Tris (Tris(hydroxymethyl-aminomethane)) (ROTH CARL, Karslsruhe, Germany; 48255.2)
- EDTA (WARCHEM, Warsaw, Poland; 0566.06)
- Acetic acid A.C.S. 80% (StanLab, Lublin, Poland; 04/56419730)
- Qubit dsDNA HS Assay Kit (ThermoFisher Scientific, Waltham, MA, USA; Q33230)
- Midori Green DNA (Nippon Genetics, Duren, Niemcy; MG10)
- DNA Gel Loading Dye 6x (ThermoFisher Scientific, Waltham, MA, USA; R0611)
- T4 DNA Ligase (5U/µl) (Promega, Madison, USA; M180A)
- Metaphor agarose (Lonza, Basel, Switzerland; 733-1211)
- Mme I (2U/µl) (New England Biolabs, Ipswich, MA, USA; R0637S)
- Agilent RNA 6000 NanoKit (Agilent, Santa Clara, CA, USA; 5067-1511)
- Agilent High Sensivity DNA Kit (Agilent, Santa Clara, CA, USA;5067-4626)
- 3% Marker C cartridges Pippin Prep (SageScience, Beverly, MA,USA; CSD3010)
- DNA ladder 100 bp (ThermoFisher Scientific, Waltham, MA, USA; SM0323)
- DNA ladder low range (ThermoFisher Scientific, Waltham, MA, USA; SM1191)

### Equipment

- Porcelain mortar
- Centrifuge
- MagJET Separation Rack (ThermoFisher Scientific, Waltham, MA, USA;
- ThermoShaker thermoblock TS-100C (Biosan, Riga, Lotwa; 010143-1301-0021)
- Deep freezer -86^°^ C
- Thermocycler
- Illumina MiSeq (Illumina, San Diego, CA, USA)
- Qubit 3 Fluorymetr (Invitrogen, Carlsbad, CA, USA; Q33216)
- Bioanalyzer 2100
- Pippin Prep (SageScience, Beverly, MA, USA; DE54107946)
- Electrophoresis system vertical gel

### Protocol

#### Total RNA isolation

The embryonic part (embryo and scutellum) of dry grains was used for isolation ot total RNA. Material for analysis was obtained from 25 seeds per biological replicate. Analysis was performed in three biological replicates for each sample. The plant material was described in detail in Puchta et al.[10].Isolations of total RNA were carried out using Trizol reagent [11] according to the following procedure:

1. Homogenize the embryos in liquid nitrogen using a porcelain mortar to produce a fine powder. Transfer up to 100 mg of this powder to a pre-chilled 2 ml tube and add 1 ml of Trizol to the ground embryos. Incubate at room temperature (23 °C) for 5 minutes.
2. Add 0.1 ml 1-bromo-3-chloropropane, incubate for 5 min. at room temperature and next centrifuge at 4 °C, in 17 700 x g, for 10 min.
3. Transfer the supernatant to a new tube, add 0,5 ml isopropanol and centrifuge at 4 °C, in 17 700 x g, for 10 min.
7. Remove the supernatant, wash the pellet with 1 ml 75% ethanol and centrifuge at 4°C, in 6 900 x g, for 5 min.
9. Remove the supernatant and dry the pellet to completely remove the ethanol.
10. Suspend the pellet in 100 µl molecular biology RNase-free water and assess the quality and integrity of RNA using a Qubit fluorometer or Agilent Bioanalyzer.

### mRNA fraction isolation

1. To prepare Dynabeads, place 200 µl Dynabeads in 2 ml tube, place on a magnetic stand for 30 seconds, remove supernatant and add 100 µl Binding Buffer.
2. Add 100 µl RNA (80 µg), mix by pipetting and incubate 5 min. at room temperature.
3. Place mixture on magnetic stand until, ’bead’ form and solution is clear.
4. Wash the beads by adding 200 µl Wash Buffer, remove supernatant and repeat step twice.
5. Dry the bead by placing the tube on magnetic stand.
6. Elute poly(A)RNA in 20 µl molecular biology RNase-free water and mix gently.
7. Heat on thermoblock for 2 min. at 65°C, 31 x g, and immediately place on a magnetic stand and transfer 20 µL of poly(A)RNA to a new tube. Critical in this step is pre-cool the tube on ice before use.

### 5’ adapter ligation

In this step it is very important to suspend the 5’SR adaptor in 120 µl molecular biology RNase-free water and store at -80 °C. Before use, denature the 5’SR adaptor at 70 °C for 2 min and place quickly on ice, use within 30 min., and do not mix. For ligation add the reaction components in the volume increased by 10% of the final reaction composition (Table 2) due to their high degree of foaming.

**Table 1.**
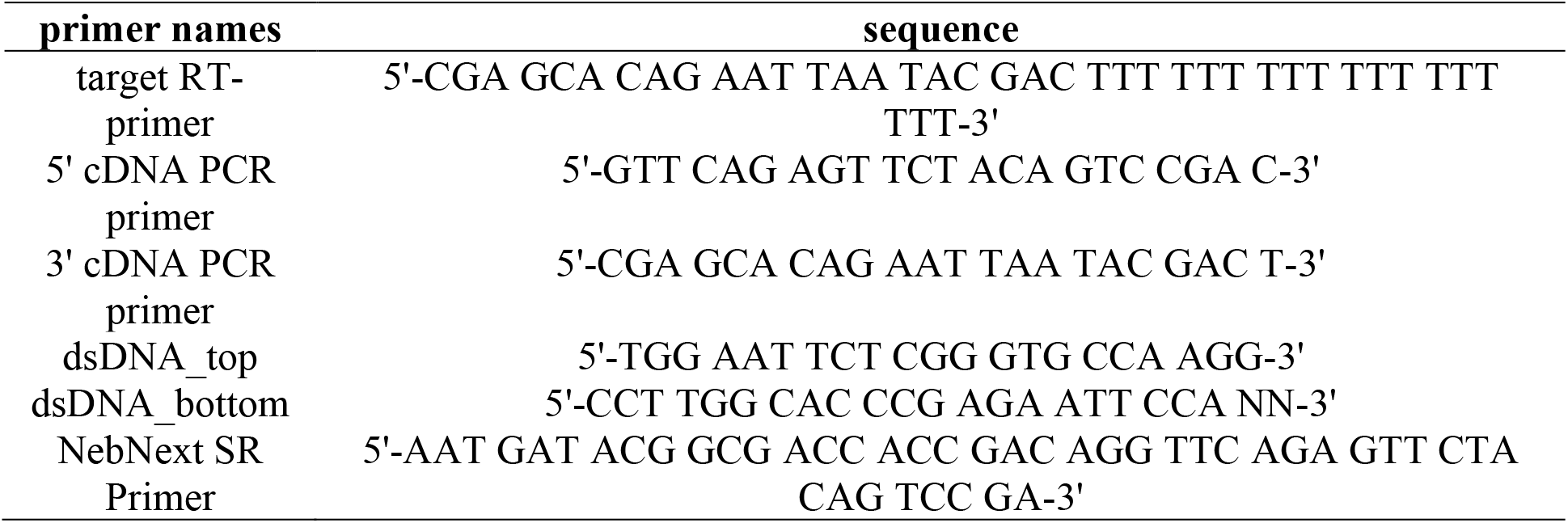
Adapter and primer sequence.

**Table 2.** Composition of the 5’ adaptor ligation reaction.

| Reagent | Volume (µl) | Final concentration |
| --- | --- | --- |
| poly(A)RNA | 15 µl | 0,3 µg |
| 5'SR adaptor (200µM) | 1 µl | 10 µM |
| 5'Ligation Reaction Buffer (10x) | 1 µl | 0,5x |
| 5'Ligation Enzyme Mix | 2,5 µl | 12,5% |
| Ribolock (40 U/µL) | 0,25 µl | 10U |
| **Total** | **20 µl** | **-** |

**Table 3.** Composition of the RT primer ligation reaction.

| Reagent | Volume (µl) | Final concentration |
| --- | --- | --- |
| RNA-target RT Primer(poly(T)) | 15 µl | 70,6 % |
| 5x Reaction Mix | 4 µl | 0,95 x |
| Maxima Enzyme Mix (200 U) | 1 µl | 200 U |
| dNTP | 1 µl | 0,5 mM |
| Ribolock (40 U/µL) | 0,25 µl | 10U |
| **Total** | **21,25 µl** | **-** |

**Tabela 4.**
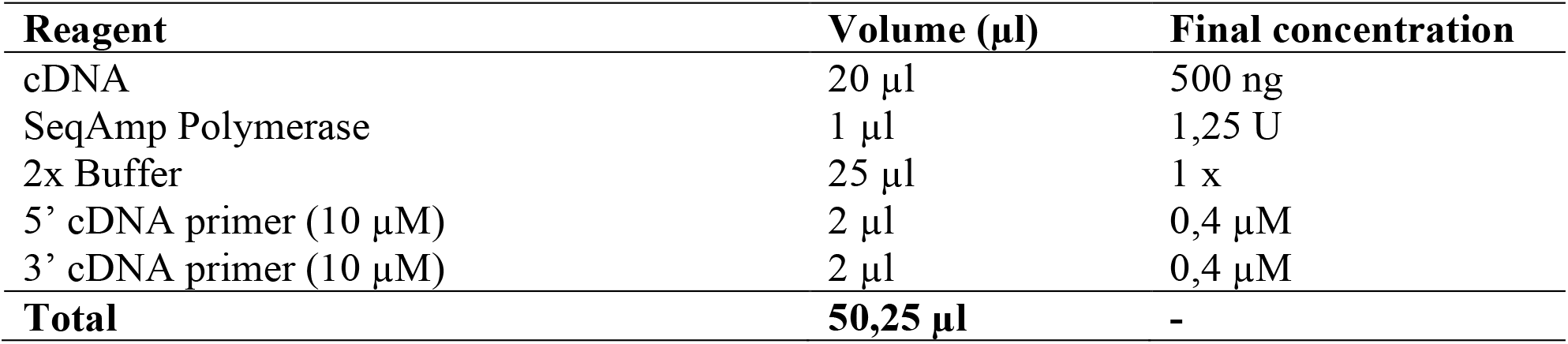
Composition of the cDNA synthesis reaction.

3. Perform the following reaction on the cDNA template: **Tabela 4.** Composition of the cDNA synthesis reaction
| Reagent | Volume (µl) | Final concentration |
| --- | --- | --- |
| cDNA | 20 µl | 500 ng |
| SeqAmp Polymerase | 1 µl | 1,25 U |
| 2x Buffer | 25 µl | 1 x |
| 5' cDNA primer (10 µM) | 2 µl | 0,4 µM |
| 3' cDNA primer (10 µM) | 2 µl | 0,4 µM |
| **Total** | **50,25 µl** | **-** |

**Table 5.** Composition of the Mme I digestion reaction.

| Reagent | Volume (µl) | Final concentration |
| --- | --- | --- |
| PCR Product | 15,8 µl | - |
| 10 xNEB 4 Buffer | 2 µl | 1 x |
| SAM | 0,2 µl |  |
| Mme I (2U/µl) | 2 µl | 4U |
| **Total** | **20 µl** | **-** |

**Tabele 6.**
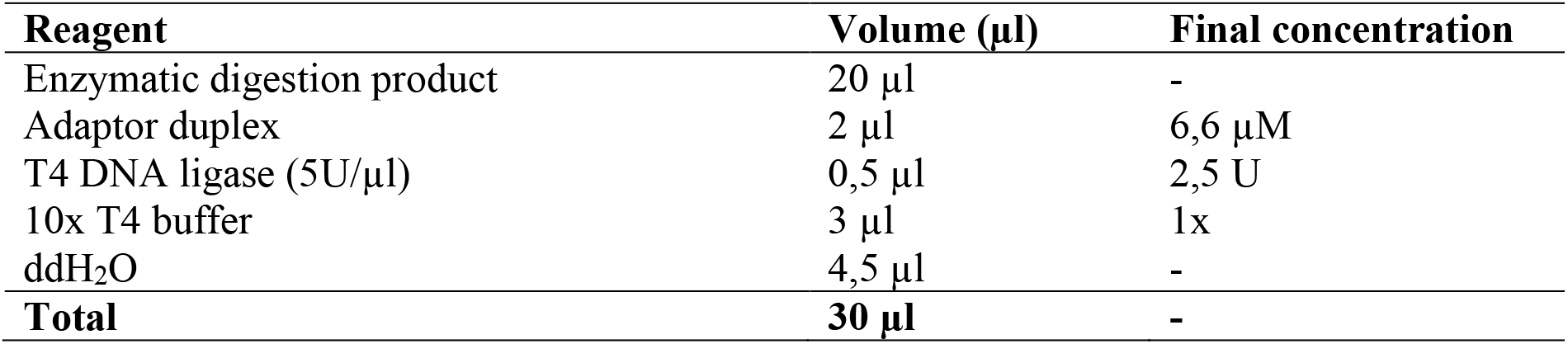
Composition of the duplex adaptor ligation reaction.

1. Add 5’ adaptor ligation reagents as follows:
2. Incubate at 37 °C for 50 min., next add 80 µl molecular biology RNase-free water and incubate at 65 °C for 10 min. then cool on ice.

### Purification using magnetic beads

1. To prepare Dynabeads, place 200 µl Dynabeads in 2 ml tube and place on a magnetic stand for 30 sec., remove supernatant and add 100 µl Binding Buffer.
2. Add 100 µl 5’SR-poly(A)RNA and mix 5 min by pipetting at room temperature.
3. Place mixture on magnetic stand until,, beads’’ form solution is clear.
4. Wash the beads by adding 200 µl Wash Buffer next remove supernatant and repeat step twice.
5. Dry the bead by placing the tube on magnetic stand.
6. Elute poly(A)RNA in 20 µl molecular biology RNase-free water and mix gently.
7. Heat on thermoblock for 2 min. at 65°C, 31 x g, and immediately place on a magnetic stand and transfer 20 µL of 5’SR-poly(A)RNA to a new tube. It is important that the tubes are pre-cooled in ice before use.

### cDNA synthesis

1. Add 1µl Target RT Primer(poly(T)) (100µM) to 14 µl 5’SR-poly(A)RNA, next incubate at 65 °C for 10 min. and quickly placed on ice.
2. Mix for cDNA synthesis was caried out according to the following composition, and incubate at 50 °C for 15 min., next at 85 °C for 5 min.
3. Perform the following reaction on the cDNA template:
4. Perform the reaction according to the following scheme:

### Clean up and quality control

In this step very important is that AmPure XP magnetic beads should be added to the DNA in a ratio of 1,6:1. AmPure XP should be kept at room temperature for 30 min. and vortex before use.

1. Add 80 µl Ampure XP to 50 µl cDNA, mix well by 10-times pipetting and incubate for 5 min. at room temperature, next place the tube on magnetic stand, and keep until the solution appears completely clear and remove supernatant.
2. Hold the tube on magnetic stand and add 200 µl fresh 70% ethanol, after 30 sec remove the ethanol, repeat step twice.
3. Air dry the beads on magnetic stand and add 17 µl molecular biology grade water to the beads and gently mix the pellet to uniformly suspend the beads.
4. Incubate the tube for 5 min at room temperature, then place the tube on magnetic stand and transfer the supernatant to a new tube. Verify the DNA quality and quantity using Qubit and Agilent 2100.

### Mme I digestation

1. Enzymatic digestion of PCR products using Mme I endonuclease should be performed in a mixture containing:
2. Incubate at 37 °C for 50 min., next at 37 °C for 50 min. and cool slowly to room temperature. It is important not to put the library on ice after digestion.

### Duplex adaptor ligation

1. Add 50 µM dsDNA top (100 µM) and 50 µM dsDNA bottom (100 µM) adaptors to a new tube and incubate the mixture at 100 °C for 5 min., then cool to 25 °C. The critical parameter is to maintain a cooling rate of 0,1 °C/ sec.
2. Prepare a ligation mix as follows and incubate 1 hour at 22 °C.

### Gel purification and precipitation

1. Prepare 4% Agarose MetaPhor gel dissolved in 1x TAE buffer and prepare3mm vertical gel
2. Add 25 µl each of samples with 5 µl loading buffer and 60 and 65 pb size ladder run on gel, run the electrophoresis at 150 V for 1,5 h.
3. To 200 ml 1 x TAE buffer add 10 ml Midori Green, place the gel in the mixture, cradle for 30 min. in the darkroom and visualize the gel at λ= 312 nm. Due to the low concentration of libraries, are visible as light smears on the gel.
4. Cut smear sections from the gel corresponding to the size of the 60 and 65 bp bands (Figure 3). Place the cut gel sections containing the library fragments into perforated tubes with sterile gauze, centrifuge at 15,000 xg for 10 min at room temperature (Figure 4).
5. Transfer the filtrate to a new tube (about 45 µl) and add 4,5 µl sodium acetate(1:10 v/v), 1 µl glycogen and 112,5 µl 99% ethanol (2,5:1v/v), mix for spin and freeze overnight at -80 °C
6. Centrifuge at 16 000 x g, for 30 min. at 4 °C, decant the supernatant and add 600 µl 70% ethanol, mix and decant the ethanol. After a centrifuge, a white to transparent precipitate should appear at the bottom of the tube, be careful not to remove the precipitate when decant the ethanol.
7. The precipitate dissolve in 20 µl of water for molecular biology, place the tube in the fridge for one hour and gently mix by tapping with fingers to dissolve the precipitate thoroughly.

**Figure 1.**
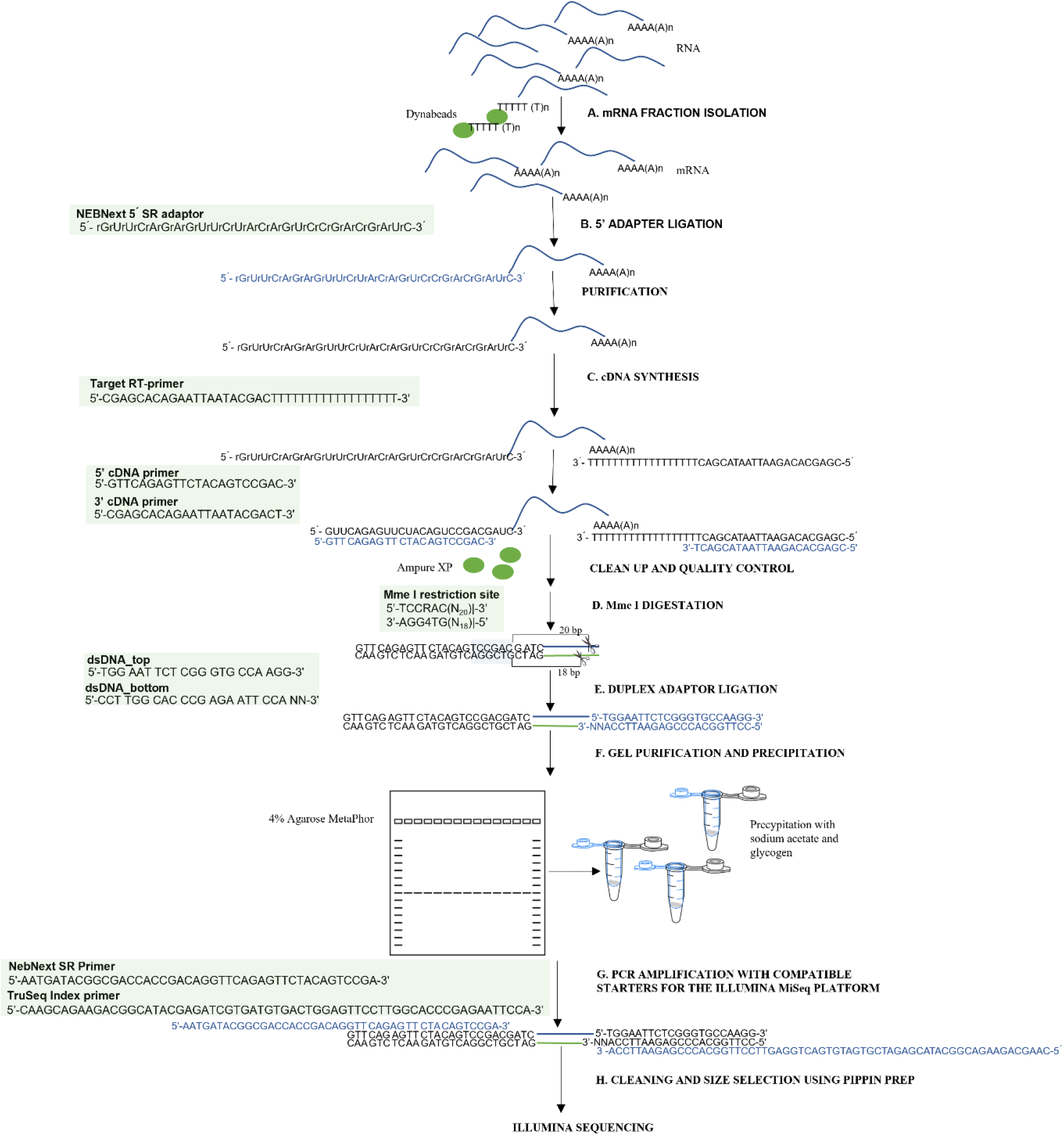
The scheme for constructing degradome library. The procedure comprises the following steps: (A.) mRNA fraction isolation, (B.) 5’ adapter ligation, (C.) cDNA synthesis, (D.) Mme I digestation, (E.) Duplex adaptor ligation, (F.) Gel purification and precipitation, (G.) PCR amplification with Illumina primers and (H.) Cleaning and size selection library using pippin prep

**Figure 2.**
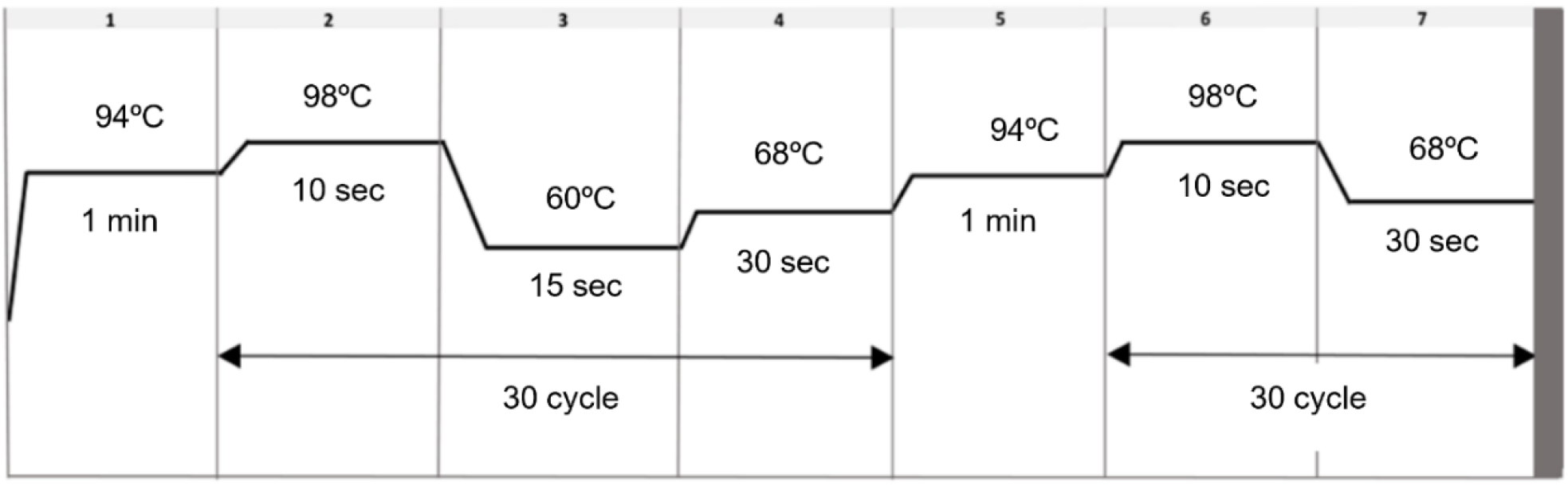
Thermal profile of the cDNA synthesis reaction

**Figure 3.**
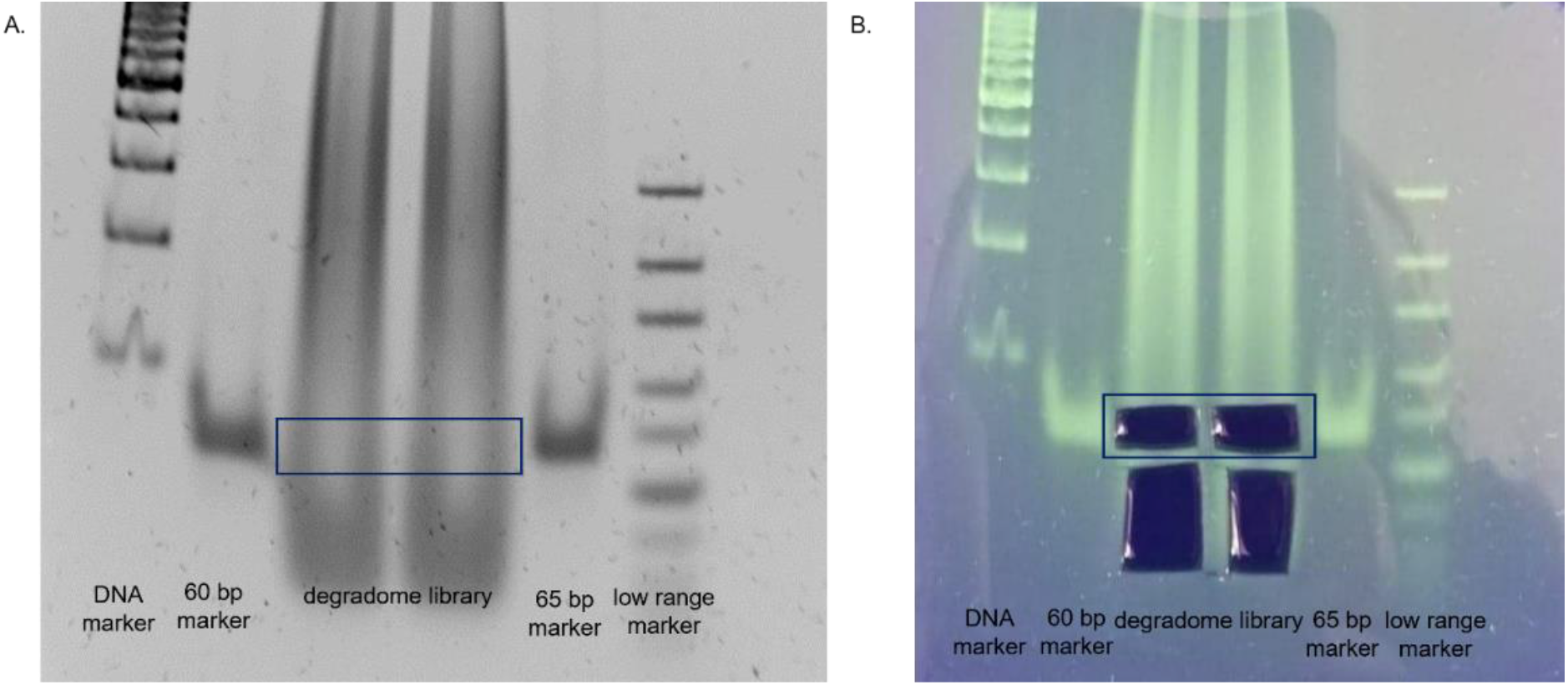
Separation of degradome libraries in 4% high-resolution MetaPhor agarose, Lonza; **A**. Electrophoretic image of the separation of libraries fragments with 60 bp and 65 bp markers; **B**. image of libraries cut extraction

**Figure 4.**
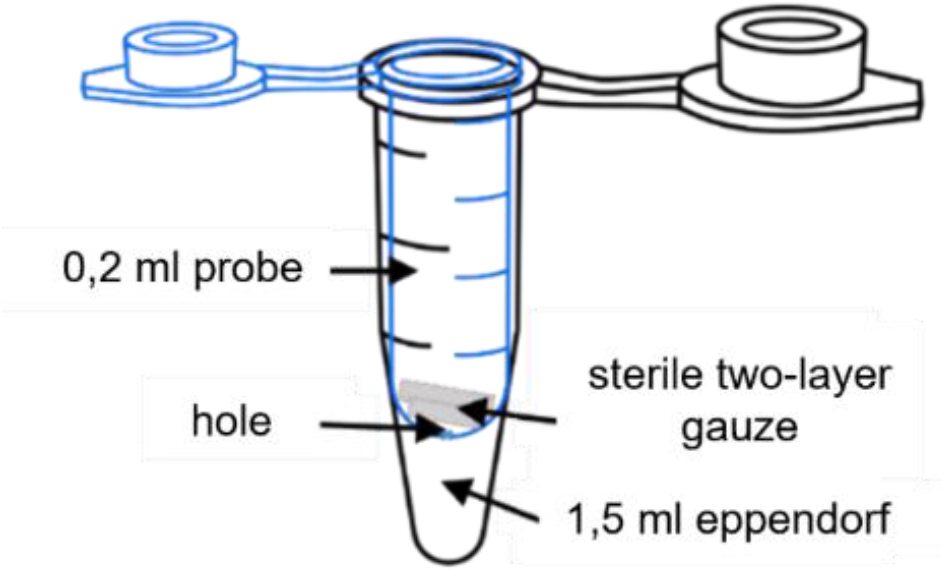
Schematic of a tube for recovery products after electrophoretic separation

### PCR amplification with starters compatible to the Illumina platform

1. Prepare a mix of the following composition (Table 7) and add to it 20 µl of purified libraries
2. Run the reaction using thermocycler according to the following thermal profile
3. Clean amplified libraries using the Pippin Prep and 3% cartridge with marker C, cleaning according to manufacturer’s recommendations. Set the parameters: start: 100; stop: 150; run time: 3h. Subsequently evaluate qualitatively and quantitatively the extracted library using a Qubit and Bioanalyzer 2100 system.

**Tabele 7.**
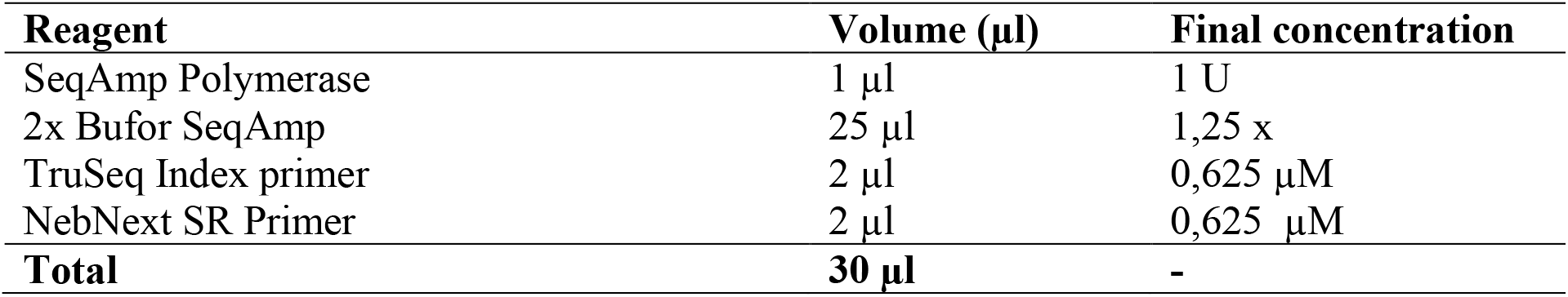
Composition of the PCR amplification reaction compatible starters for the Illumina platform.

**Table 88.** Concentration of 60, 65 bp libraries extracted from high-resolution 4% metaphor agarose and 12% polyacrylamide using one of three purification methods (1. freezing fragments with glycogen added; 2. centrifuge on gauze; 3. centrifuge on gauze with glycogen added).

| library | methods of extraction | amount applied on gel [ng] | Qubit measuring [ng] | percent of recovery [%] |
| --- | --- | --- | --- | --- |
| MetaPhor Agarose 4% |  |  |  |  |
| degr60 | 1 | 298 | 3,3 | 4,46 |
|  | 2 | 334 | 21,2 | 6,35 |
|  | 3 | 320 | 59 | 18,4 |
| degr65 | 1 | 232 | 9,3 | 9,3 |
|  | 2 | 206 | 12,3 | 12,3 |
|  | 3 | 244 | 38,8 | 15,9 |
| Poliacrylamide 12% |  |  |  |  |
| degr60 | 1 | 298 | 0,2 | 0,06 |
|  | 2 | 334 | 1,04 | 0,3 |
|  | 3 | 320 | 12,2 | 3,81 |
| degr65 | 1 | 232 | 1,8 | 0,8 |
|  | 2 | 206 | 6,9 | 3,35 |
|  | 3 | 244 | 4,3 | 1,76 |

### Illumina Miseq platform sequencing

1. Add 58,9 µl water for molecular biology grade to 2 µl library (2 nM library)
2. Add 5 µl 0,2 N NaOH to 5 µl 2 nM library, next vortex, centrifuge at 280 x g, for 1 min. and incubate for 5 min. at room temperature, then add 990 µl pre-cooled HT1 Hybridization Buffer and not vortex. The final denatured library contains 20 pM.
3. Add 3 µl 10 mM Tris-HCl to 2 µl 10 nM PhiX control, than ad 5 µl 0,2 N NaOH next vortex, centrifuge at 280 x g, for 1 min. and incubate for 5 min. at room temperature, then add 990 µl pre-cooled HT1 Hybridization Buffer and not vortex. The final denatured PhiX control library contains 12,5 pM.
4. Add 225 µl HT1 Hybridization Buffer to 375 µl 20 pM PhiX and mix by inverting.
5. Add 594 µl denatured library to 6 µl denatured and diluted PhiX and apply 600 µl library on Illumina cartridge and not vortex.

Perform sequencing e.g. using Reagent Kit v3 (50 cycles) (Illumina) on Illumina MiSeq. Read length 36 bp, sequencing from a single end (single end).

## Results and discussion

To prepare the protocol, a number of optimizations and verifications were performed. See Appendix 1 for a detailed description. The protocol has two notable innovations. Firstly, it allows for more efficient extraction of libraries prior to binding of Illumina adapters, thereby enhancing the overall efficiency of the method. Secondly, it employs automated electrophoresis as the final purification technique for libraries, which ensures superior quality of the final library. Tests were conducted to identify the most effective methodology for visualizing and extracting libraries from gels in order to enhance the overall library construction process. The fragments must be 60–65 bp in length, precluding purification of the samples via both automatic electrophoresis and magnetic beads. The components and conditions of the individual steps in the preparation of the libraries have been optimized for challenging materials such as grains of long-term stored cereals that are rich in reserve substances, e.g., sugars. To this end, primers designed for the ADP-ribosylation factor gene (NCBI accession number 838957) were used to amplify fragments of 60 and 65 base pairs in length, which were then utilized as size markers to facilitate the visualization and extraction of degradome libraries in subsequent steps (Figure 6A).

**Figure 5.**
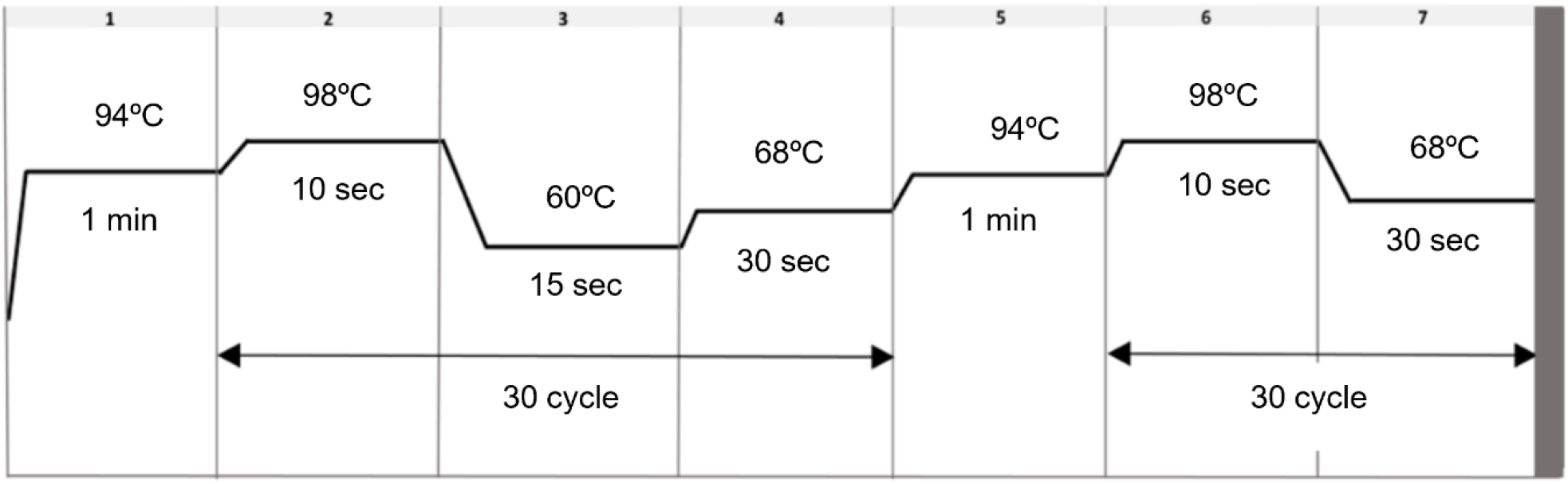
Thermal profile of the cDNA PCR amplification reaction compatible starters for the Illumina platform

**Figure 6.**
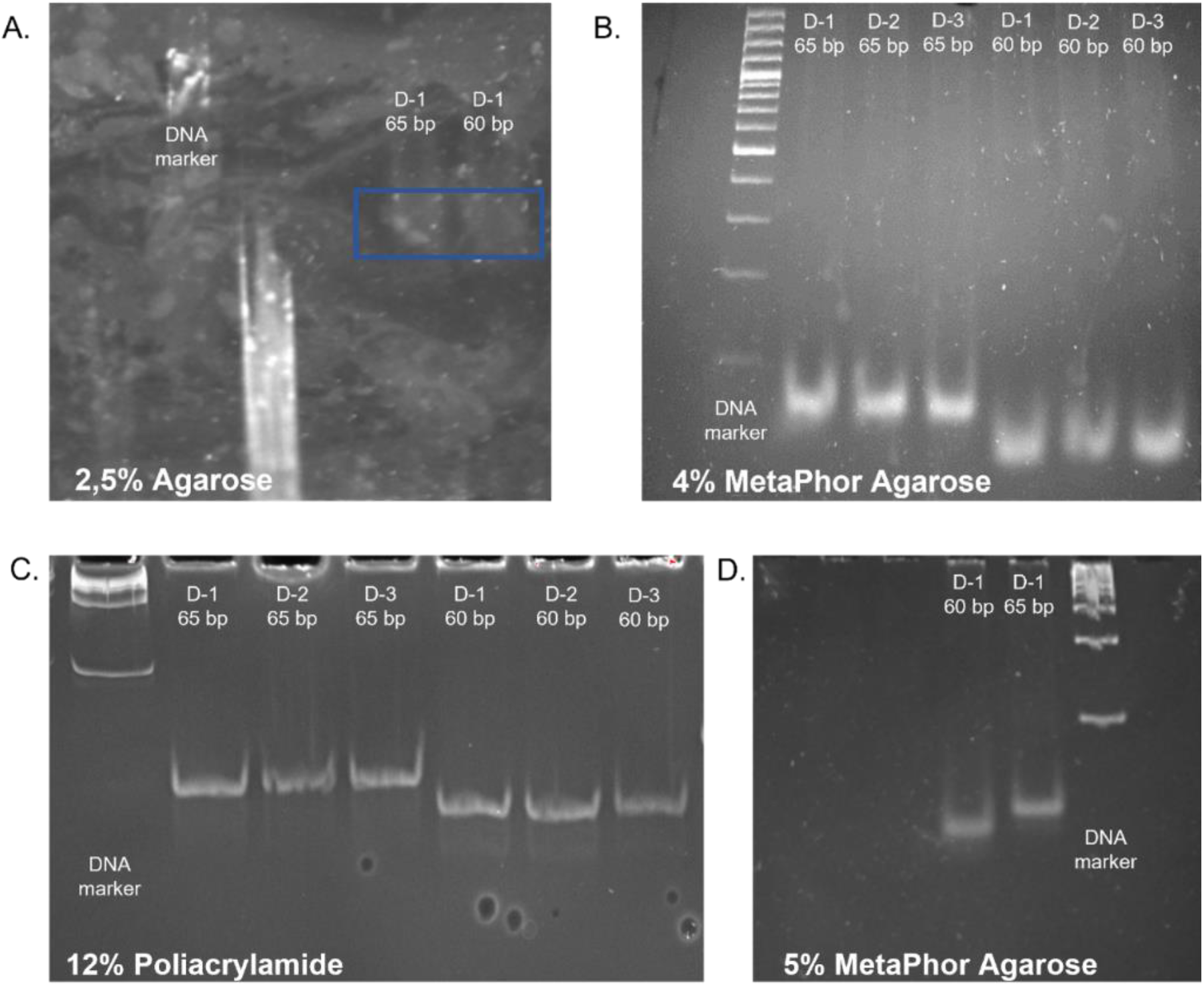
A qualitative assessment of the separation of 60, 65 bp samples, the separation was performed in: A. 2.5% agarose gel (samples were mark by box); B. 4% MetaPhor high-resolution agarose, Lonza; C. 12% polyacrylamide gel; D. 5% high-resolution MetaPhor agarose (D-1; D-2; D-3-technical replicates).

To achieve enhanced separation, high-resolution MetaPhor agarose at concentrations of 5% and 4% were evaluated (Figure 6B, D) [12]. Given the challenges associated with dissolving agarose and preparing vertical gels at a 5% concentration, agarose at a 4% concentration was selected for subsequent testing. To facilitate comparison, sample separation was also performed in a 12% polyacrylamide gel. However, despite the high-resolution images of the products obtained upon visualization, the recovered libraries had very low concentrations (Figure 1C). The 4% high-resolution MetaPhor agarose gel was selected as the optimal gel for separations and visualization of degradome libraries based on the results of the electrophoresis analysis and the final product concentration. The utilization of high-resolution agarose facilitated the observation of size marker bands. The employment of high-resolution agarose for the separation of degradome libraries and the utilization of dedicated size markers enabled the observation of a library smear and the precise excision of a gel section containing fragments within the 60-65 bp range. This is in contrast to the methodology employed by other research teams, who excised fragments based solely on the size marker [13].

The 60 bp and 65 bp PCR products were cut from the gel and DNA fragments were purified using common laboratory consumables in place of specialized, expensive cellulose acetate or nylon membrane centrifuge tube filters. Therefore, a 1.5ml Eppendorf tube was used with a 0.2ml tube placed inside (Figure 4). A preparation needle was used to make a hole in the bottom of the 0.2 ml tube. On the bottom of the smaller tube, two layers of sterile autoclavable gauze were placed. A solution containing sodium acetate, ethanol and glycogen was then added to each filtrate to precipitate the DNA.

The solution was enriched by adding glycogen due to the low concentration of fragments in the obtained filtrates. Glycogen is a polysaccharide that can effectively co-precipitate with DNA in the presence of alcohol, such as ethanol or isopropanol. It aggregates with DNA, increasing the weight of the molecules and facilitating precipitation [14]. For samples with low concentrations of DNA, the addition of glycogen significantly increases the efficiency of the precipitation procedure. Low-concentration DNA can be difficult to precipitate, but glycogen helps form larger, more easily centrifuged aggregates. Thus, even very small amounts of DNA can be recovered which might be left in the supernatant without glycogen [15]. Due to their low efficiency for fragments less than 70 bp in length and low initial concentration, none of the commercially available agarose gel elution kits were used. Three methods were tested for eluting and precipitating degradome library fragments after agarose and polyacrylamide gel electrophoresis i.e. freeze and squeeze method in a glycogen-containing solution, centrifuged in gauze-containing tube with standard eluent without glycogen and centrifuged in gauze-containing tube with solution enriched with glycogen. The concentration of the DNA fragments was measured prior to electrophoresis and after elution from the gel using a Qubit fluorimeter and a dsDNA HS Assay Kit. Based on the results, the highest sample recovery of almost 20% was obtained using the third method from agarose gel. The efficiency of elution from the polyacrylamide gel was undoubtedly influenced by the type of buffer used. It is standard practice to use high ionic strength buffers containing EDTA in their composition for elution [16, 17]. However, the presence of EDTA in samples undergoing NGS sequencing is unfavorable due to inhibition of MG^2+^ ion-dependent enzymes, such as ligase and polymerase. This can contribute to incomplete or inefficient amplification and ligation, resulting in lower sequencing yield and quality [18, 19]. Due to the low concentration of DNA fragments, EDTA could not be removed by further purification steps.

The protocol for generating degradome libraries was validated using genetic engineering tools. The final fragments of the libraries were cloned into the pGEM-T Easy Vector System vector. The vector was electroporated into competent Escherichia coli DH5α bacteria. The pGEM-T Easy Vector System vector makes use of a plasmid with a length of 3015 bp. The insertion site in the vector is located in the lacZ gene, which encodes the ß-galactosidase subunit. Clones containing the correct insertion were selected by adding X-Gal and ampicillin to the medium. Alkaline lysis was performed to obtain plasmid DNA. For evaluation of the alkaline lysis process, electrophoresis was performed on a 1.5% agarose gel on which 5 µg of samples were loaded (Figure 7).

**Figure 7.**
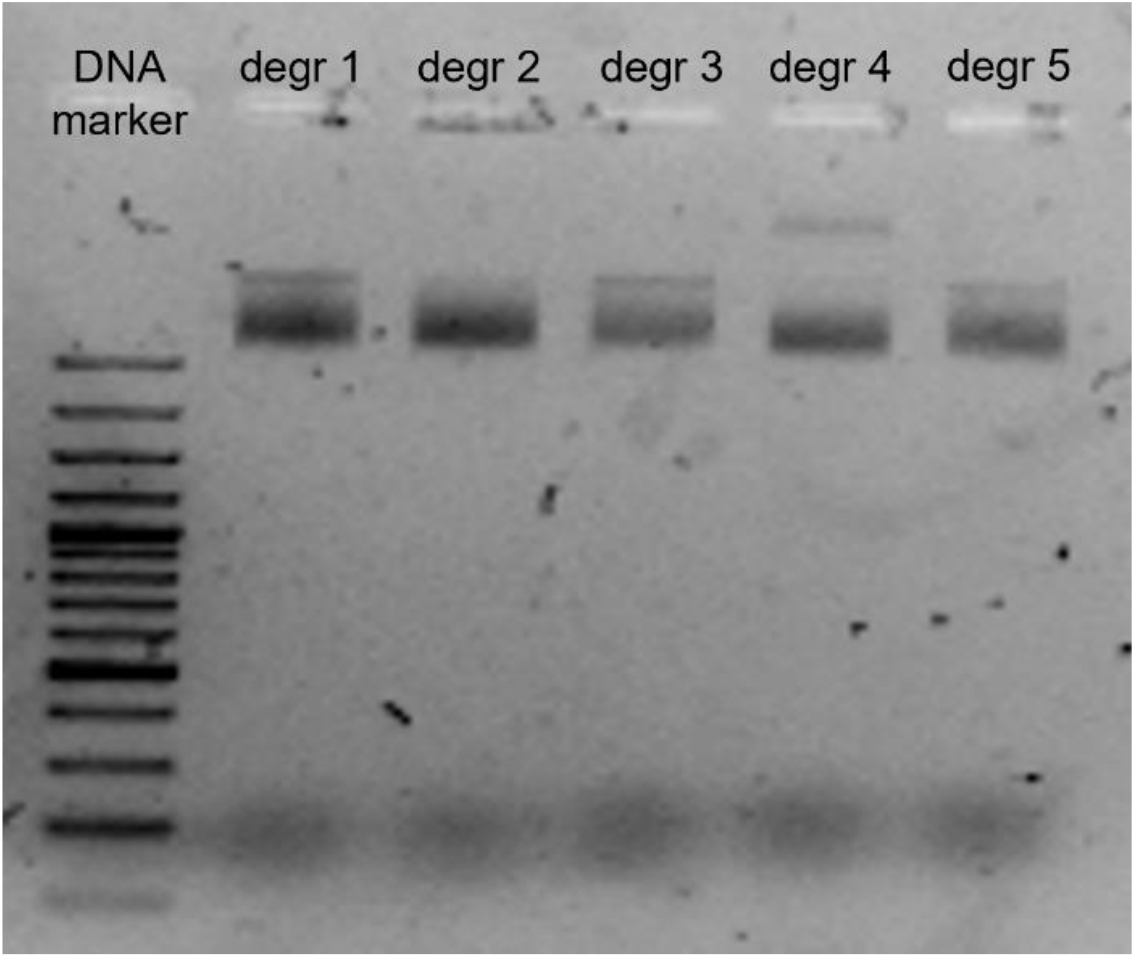
Electrophoretic image of plasmid DNA from *E*.*coli* (plasmid DNA contained a degradome library insert).

Sanger sequencing was performed on the fragments containing the integrated degradome library fragment. The sequences obtained confirmed the correct binding of all adaptors to the fragment obtained after restriction with the enzyme Mme I (Figure 8)

**Figure 8.**
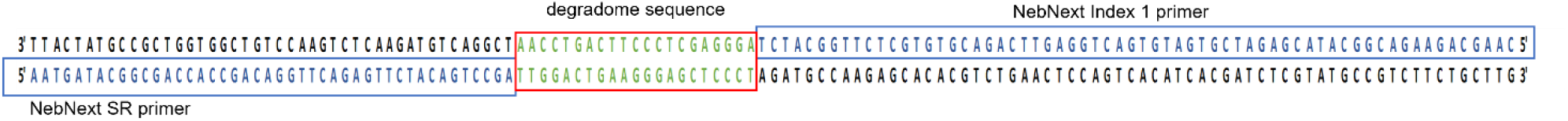
The degradome library sequence obtained by Sanger sequencing.

During the analysis, libraries of 130-150 bp were confirmed their size using Bioanalyzer 2100 automatic electrophoresis (Figure 9). Following the steps described in protocol, constructed library with produced 25 mln. single reads using MiSeq Reagent Kit v 3. As a result of sequencing, 7791,680, 7858,994 and 6797,471 raw reads were obtained. The high quality of the reads (Figure 10) was confirmed by evaluation of the reads obtained during sequencing using FastQC [20]. The parameter indicating the quality of the sequencing (QC) was 98.5%. The qualitative analysis of the raw reads indicated their high quality.

**Figure 9.**
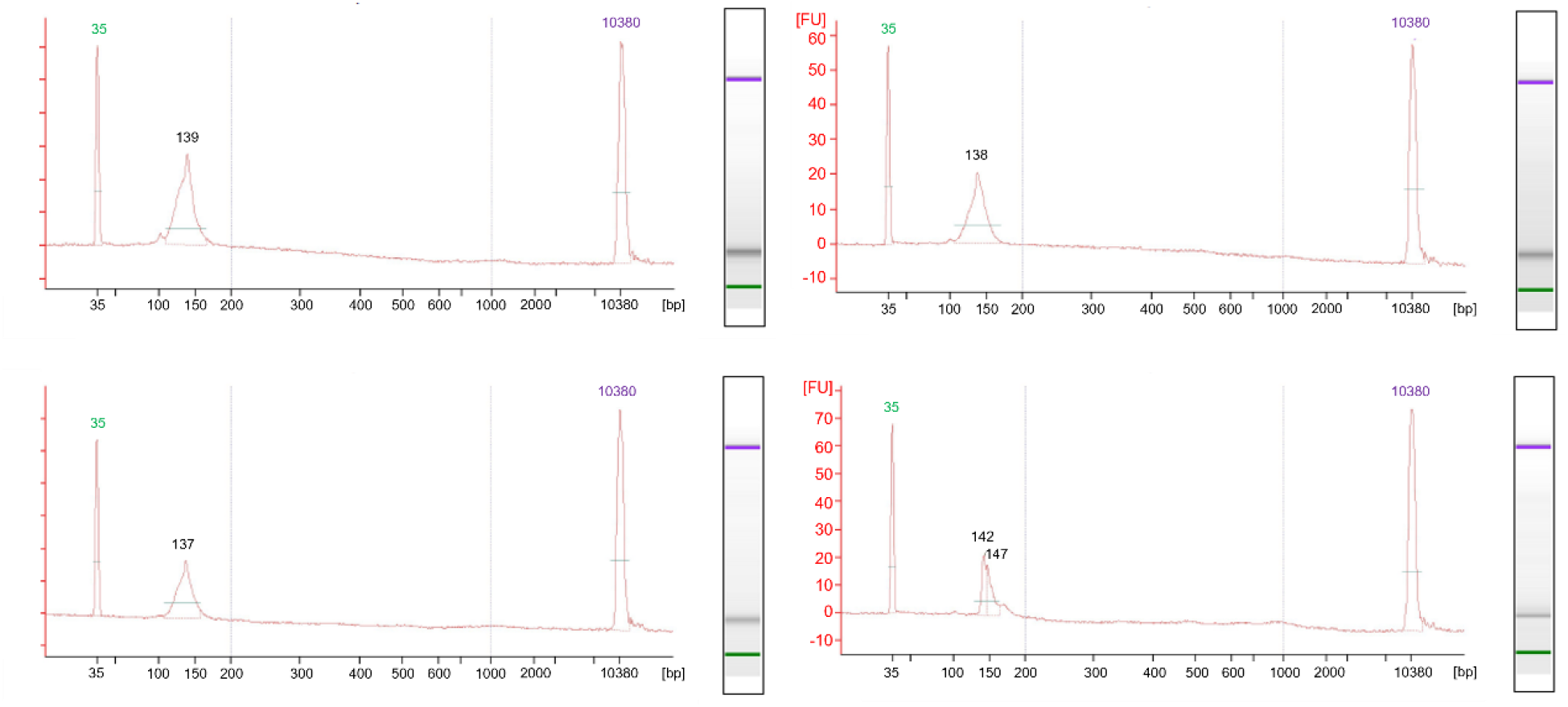
Result of degradome library construction described in this study, using 2100 Bioanalyzer High Sensivity kit (Agilent). Electrophoregrams described library size.

**Figure 10.**
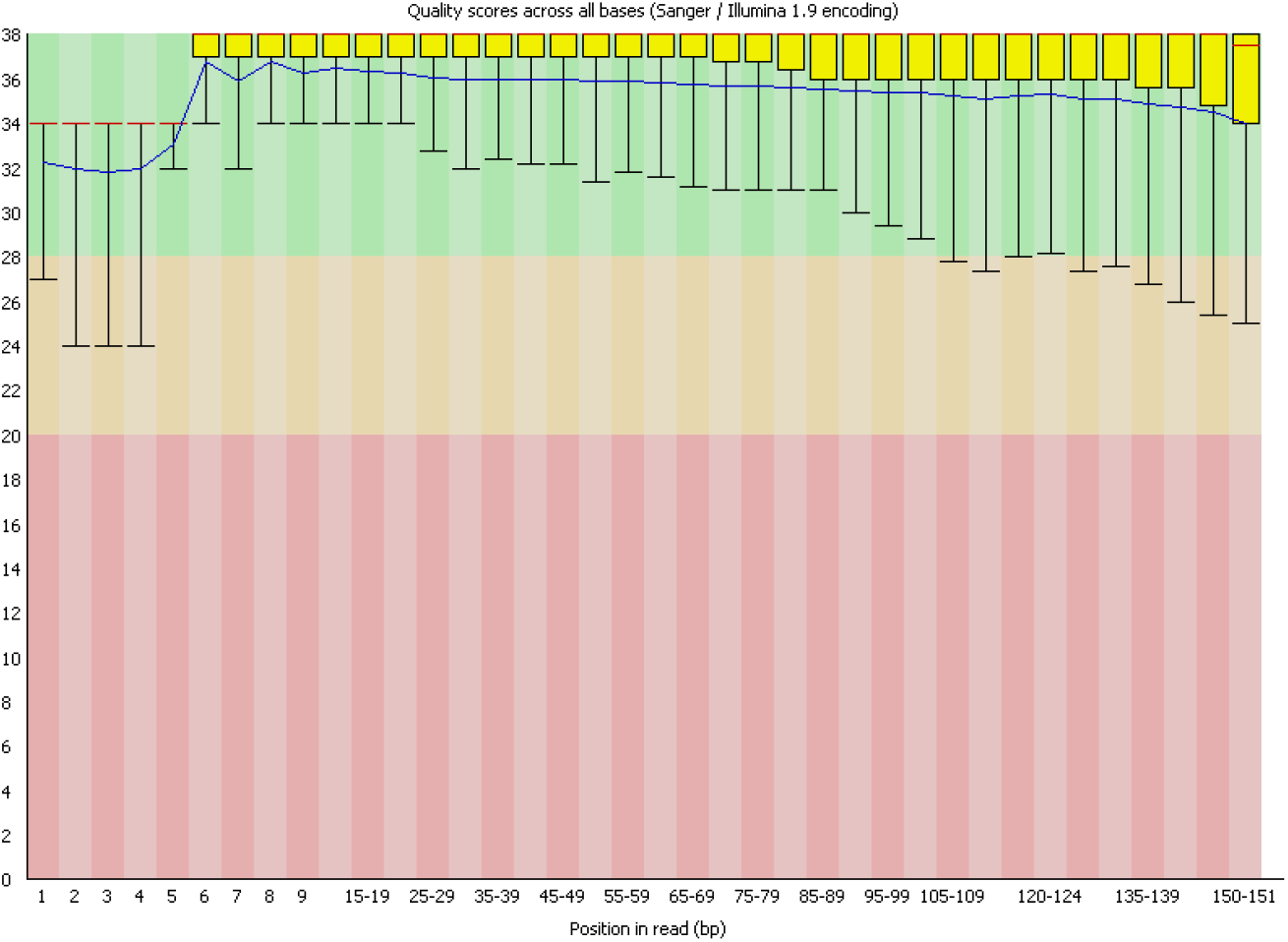
Qualitative analysis of degradom libraries, performed with the FastQC program.

The developed protocol has been used to verify targets for miRNAs detected in dry seeds and during the first 24 hours of germination of barley seeds after long-term storage. Further research aims to identify the relationship between germination capacity and miRNAs and seed aging processes.

The degradome analyses significantly improved the identification of miRNA target sequences in radish, rice and cotton [21]. The preparation of high quality degradome libraries allowed the verification of false positives from *in silico* analyses [22].

The developed protocol allowed the construction and sequencing of degradome libraries from very difficult material, such as barley grains after long-term storage, which were characterized by very uneven degradation of various RNA fractions and low RIN [23]. Moreover, the protocol does not require the purchase of additional reagents and is based on the ingredients included in the kit for sRNA analysis, thus significantly reducing the cost of library construction. The use of additional dedicated size markers allows for the precise excision of a band containing the appropriate length of fragments from the gel. In comparison to previously used methods based on polyacrylamide gels, the optimization of the purification method of degradom-seq libraries allowed an increase in the yield of fragments obtained [7]. It is noteworthy that the time requirement for the complete library preparation protocol does not exceed three days, which is also a significant time saving.

## Supporting information

Appendix 1. Validation of library design by vector cloning and Sanger sequencing

## Acknowledgements

The development of the method was supported by NCN grant Preludium 18 (2019/35/N/NZ9/01046)

## Author information

Present address: Plant Breeding and Acclimatization Institute, Radzików, 05-870 Błonie, Poland

### Contributions

Conceptualization, M.P.J and J.G. M.B; methodology, M.P.J and J.G.; validation, J.G. and M.P.J; formal analysis, J.G. and M.P.J; investigation, M.P.J and J.G., M.B; resources, M.B.; writing—original draft preparation, M.P.J; writing—review and editing, M.P.J, J.G. and M.B.; visualization, M.P.J; supervision, J.G. and M.B. All authors have read and agreed to the published version of the manuscript.

## Corresponding author

Correspondence to Marta Puchta-Jasińska

