## Appendix 1. Validation of library design by vector cloning and Sanger sequencing for "Improved Degradome Sequencing Protocol via Reagent Recycling from sRNAseq Library Preparations"

Library cloning was performed using Escherichia coli DH5α competent bacteria and the pGEM-T Easy Vector System vector.

In the vector, the insertion site is located in the lacZ gene, encoding the ß-galactosidase subunit. Selection of clones with the correct insertion is performed by adding X-Gal and ampicillin to the substrate.

A-tilling (addition of adenine at the 3' end) was performed first, for which a reaction was prepared with the following composition (Table 1) and a total volume of 3 µl, which was incubated at 70 °C for 30 minutes.

**Table 1. Composition of the A-tilling reaction**

| Reagent | Volume |
|---|---|
| library degradome-Seq | 3,2 ng |
| 2mM dATP | 0,2 mM |
| 10x DreamTaq buffer | 1x |
| 5U/µl DNA DreamTaq polimerase | 1,5 U |
| total | 3 µl |

The ligation of the degradome libraries to the plasmid was then carried out. For this purpose, a reaction was prepared with the composition shown in Table 2.

**Table 2. Composition of the ligation library to plasmid reaction**

| Reagent | Volume |
|---|---|
| A-tiling reaction | 30% |
| pGEM T-easy plasmide | 10% |
| T4 ligase | 3U |
| 2x buffer | 1x |
| total | 10 µl |

Incubated at 16 °C overnight.

Plasmids obtained by electroporation were transformed into competent cells of E.coli, strain DH5α. To adopt fragments of the degradome libraries, the bacterial cells were exposed to short electrical pulses of 2.5 kV. To recover from the electrical shock disintegrating the stability of the cell membranes, bacteria were incubated in SOC medium in a thermoblock at 37 °C for 1 h at 250 rpm. SOC medium at pH 7 was prepared according to the composition in Table 3.

**Table 3.** Composition SOC medium

| Reagent | Concentration |
|---|---|
| tryptone | 2% |
| yeast extract | 0,5% |
| NaCl | 8,6 mm |
| $MgCl_2$ | 10 mM |
| glucose | 20 mM |
| KCl | 2,5 mM |
| bacteriological agar | 1% |

200 µl of bacteria were inoculated onto LB solid medium (Table 4) with ampicillin, pH 7, and the addition of selection factors 0.1mM X-Gal and 6.9 µM IPTG, and incubated at 37 °C overnight.

**Tabele 1.** Composition LB  medium with ampiciline and X-Gal, IPTG

| Reagent | Concentration |
|---|---|
| tryptone | 1% |
| yeast extract | 0,5% |
| NaCl | 172 mM |
| KCl | 13 mM |
| Bacteriological agar | 1% |
| ampicyline | 286 mM |

Reduction cultures were made from white bacterial colonies on LB media with ampicillin and incubated at 37 °C overnight. Selected bacterial colonies were transferred to 3 ml of LB liquid media with ampicillin (Table 5). Bacterial culture was conducted at 37 °C overnight at 250 rpm.

**Tabele** 5. Composition LB medium

| Reagent | Concentration |
|---|---|
| tryptone | 1% |
| yeast extract | 0,5% |
| NaCl | 172 mM |
| KCl | 13 mM |
| ampicyline | 286 mM |

Next, alkaline lysis was performed to obtain plasmid DNA from the bacterial cultures. For this, the bacterial culture was cooled on ice and centrifuged at 22 °C for 1 min at 13 000 rpm. The resulting supernatant was removed, the pellet was resuspended in GTE buffer, pH 8 (Table 6), and 5 µl RNase was added, mixed gently and incubated at 22 °C for 5 min.

**Tabele 2.** Composition GTE buffer

| Reagent | Concentration |
|---|---|
| glukose | 50 mM |
| Tris HCl | 25 mM |
| EDTA | 10 mM |

Subsequently, 200 µl of a 1% solution of SDS with 0.2 M NaOH was added. Gently mixed by rotating the tubes and placed on ice. After 5 minutes of incubation, 150 µl of cold solution III with the composition shown in Table 7 was added to the homogeneous, transparent lysate.

**Tabele 7.** Composition III lise buffer

| Reagent | Concentration |
|---|---|
| 5 M KoAc | 0,3 mM |
| cold acetic acid 99,5% | 11,5 mM |

The mixture was then shaken vigorously and left on ice for 10 minutes. The mixture was centrifuged at 22 °C for 10 minutes at 13 000 rpm. The supernatant was removed and the pellet was washed with 500 µl of 70% ethanol. In the next step, the pellet was centrifuged at 22 °C for 10 minutes at 13,000 rpm, dried and dissolved in 30 µl of DNAz and RNAz free water for one hour in the refrigerator.

The PCR reaction was then performed with two pairs of primers, T7 and SP6, and M13 (Table 8) using the HGC PCR Mix Plus kit with the following composition (Table 9) and thermal profile (Figure 1).

**Tabele 8.** Primer sequences

| Primer | Sequence |
|---|---|
| T7 | 5’ TAATACGACTCACTATAGGG 3’ |
| SP6 | 5’ ATTTAGGTGACACTATAGAA 3’ |
| M13_R | 5’ GTAAAACGACGGCCAGT 3’ |
| M13_F | 5’ CAGGAAACAGCTATGAC 3’ |

**Tabele 9.** Composition PCR Mix Plus HGC

| Reagent | Concentration |
|---|---|
| PCR Mix Plus HGC 2x | 1x |
| forward primer | 0,4 µM |
| revers primer | 0,4 µM |
| DNA template | 50 ng |
| $H_2O$ | to final volume |

| | |
|---|---|
| Final volume | 10 µl |

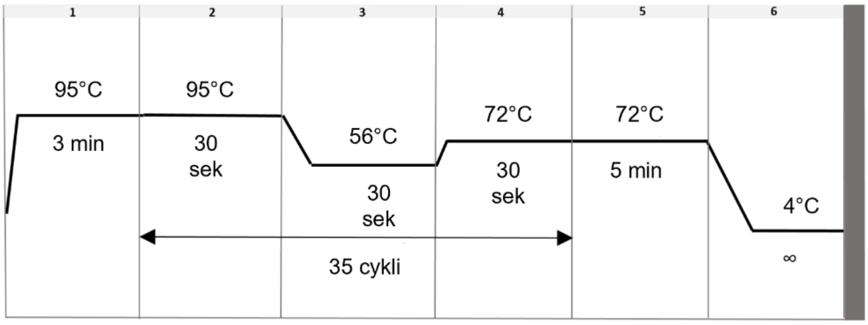


**Figure 1.** Thermal profile PCR HGC reaction

Reaction products were separated electrophoretically in a 1.5% agarose gel and visualised in the presence of Midori Green dye under UV light (λ=312 nm) using a Proxima C16 imaging system. PCR products were purified from post-reaction components using Eppic Fast. Sample concentrations were then measured using a Qubit fluorimeter with the Qubit dsDNA HS Assay Kit and diluted to 5 ng/µl according to the manufacturer's protocol for sequencing reactions of 100-200 pz fragments using the BrightDye Terminator Cycle Sequencing Kit. Sequencing PCR was performed according to the composition of the reaction mixture in Table 16 and the thermal profile according to Figure 2.

**Tabele 10.** Composition Sanger sequencing with BrightDye Terminator Cycle Sequencing Kit (MCLAB)

| **Reagent** | **Volume** |
|---|---|
| Bright Dye | 5% |
| 5x sequencing Buffer | 0,875x |
| primer | 3,2 pM |
| DNA template | 5 ng |
| $H_2O$ | to final volume |
| final volume | 10 µl |

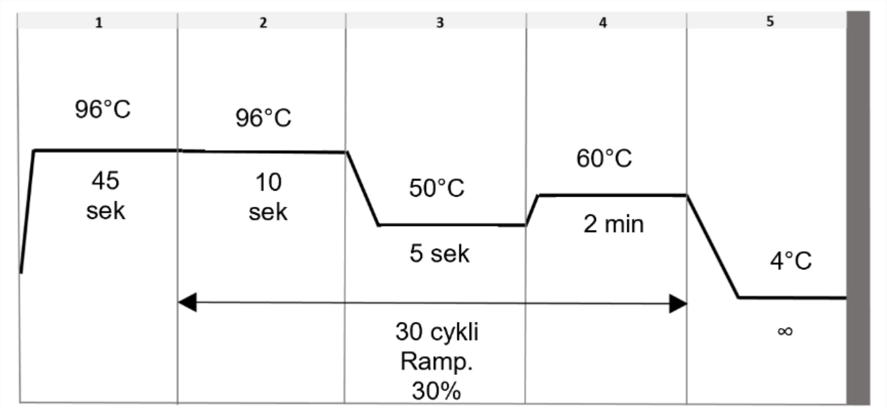


**Figure 2.** Thermal profile Sanger sequencing PCR

PCR products were purified with the ExTerminator kit. Purified products were denatured at 80 °C for 3 min and then sequenced on a 3130 XL Genetic Analyzer capillary sequencer using 36 cm capillaries and NanoPOP-7 polymer.
